# Comprehensive lipidomics of tissue macrophages reveal LTE4–driven eosinophil survival

**DOI:** 10.64898/2026.06.11.731529

**Authors:** Magdalena A. Czubala, Patricia Rodrigues, Natacha Ipseiz, Marcela Rosas, Sandra Dimonte, Iestyn Pope, Christine Hinz, Dina Fathalla, Jorge Alvarez-Jarreta, Victoria J. Tyrrell, Wolfgang Langbein, Paola Borri, Robert Andrews, Valerie B. O’Donnell, Philip R. Taylor

## Abstract

Tissue-resident peritoneal macrophages (pMФ) programmed by GATA6 are essential players regulating immunity and tissue balance in the peritoneal cavity. How GATA6 regulates global lipid metabolism is currently unknown. Addressing this, in myeloid-restricted *Gata6*-deficient (G*ata6*-KO^mye^) mice, significant changes to the pMФ lipidome were found. First, Bodipy staining and anti-Stokes Raman scattering (CARS) microscopy demonstrated significant intracellular lipid accumulation in lipid droplets in G*ata6*-KO^mye^. Untargeted and targeted lipidomics revealed this to result from increased levels of multiple sphingolipid (SL) molecular species, including sphingomyelins, ceramides, and glycosphingolipids, along with upregulation of the cysteinyl leukotriene (CysLTs) pathway at both lipidomic and transcriptional levels. Evidencing a functional role for the lipidomic phenotype, G*ata6*-KO^mye^ showed significant eosinophil accumulation, associated with decreased apoptosis which was exclusively driven by CysLT signalling. In summary, GATA6 is demonstrated as a regulator of sphingolipid accumulation and CysLT generation in pMФ, with secondary impacts on associated leukocytes through regulation of transcellular CysLT signaling.

## Introduction

Tissue resident macrophages are locally programmed by their microenvironment and specific transcription factors, as exemplified by the established role of GATA6 in driving specialization of peritoneal macrophages (pMФ)^1–7^ to fulfil their broad immune and homeostatic functions^8^. GATA6 is a zinc finger transcription factor, induced by retinoic acid, which is present in the peritoneal cavity. GATA6-expressing pMФ maintain the peritoneal cavity microenvironment, regulate inflammation and contribute to repair of organs locally^9^. Mice with a myeloid-specific deficiency of GATA6 (*Gata6*-KO^mye^) have lost their pMФ tissue-specific transcriptional programming and show abnormalities in proliferative renewal, regulation of B-1 cell-mediated gut IgA production, and marked increases in peritoneal eosinophil and F4/80^low^ macrophage populations^6, 7^.

Little is known about the tole of lipids in regulating pMФ biology, specifically as a consequence of GATA6-mediated cell programming. However, lipid metabolism is central to macrophage function, for example, effective cholesterol clearance is required for myelin phagocytosis by microglia, 12/15-lipoxygenase (LOX) in resident MФ directs clearance of apoptotic cells during inflammation, and the prostanoid/interleukin(IL)-10 pathway regulates IL-1β release from pMФ^10–12^. Beyond pMФ, dysregulated lipid metabolism in other macrophage populations is associated with dysfunction in many cellular processes, such as their response to toll-like receptors, while lipid accumulation and oxidation in macrophages is well known in numerous diseases, such as obesity, and atherosclerosis^12, 13^. Considering this, in this study, the ability of GATA6 programming to modulate lipid metabolism and signaling will be investigated in a myeloid-restricted *Gata6*-deficient (G*ata6*-KO^mye^) mouse strain. First, global changes will be screened for using lipid imaging, CARS microscopy, and untargeted lipidomics. Following this, targeted lipidomics will validate significantly changed lipid categories, and their functional significance will be investigated using *in vivo* models.

Using these approaches, we will demonstrate that the pMФ sphingolipidome is highly dependent on GATA6-mediated differentiation, and dysregulated CysLT signaling in the absence of the transcription factor promotes local eosinophil survival and accumulation.

## Materials and Methods

### Animals

All animal experiments were performed in accordance with the United Kingdom Home Office Animal (Scientific procedures) act of 1986. *Gata6*-WT and -KO^mye^ mice were generated from Lysozyme M Cre-recombinase ‘knock-in’ congenic mice on the C57BL/6 background (‘*Lyz2^Cre^*, B6.129P2-*Lyz2^tm^*^1^(cre)*^Ifo^/J*, RRID:IMSR_JAX:004781) and conditional ‘floxed’ GATA6 deficient mice (‘*Gata6^Fl^*, *Gata6^tm^*^2^*^.1Sad/J^, RRID:IMSR_JAX:008196*), obtained from the Jackson Laboratory and bred in our animal facilities as previously described^6^. Homozygous 12/15-LO-deficient mice B6.129S2-Alox15^tm1Fun^/J (Strain#:4155, RRID:IMSR_JAX:004155) and C57BL/6J (Strain#: 000664,RRID:IMSR_JAX:000664) mice were obtained from Jackson Laboratory.

All mice were used at 8-12 weeks of age, unless stated otherwise in selected experiments. Unless stated otherwise, data points represent pooled samples from multiple female mice, using age- and sex-matched groups. All animals were given free access to water and standard chow and were housed in filter top or conventional cages (12 h light/dark cycle, temperature 21°C-23°C, humidity 30-60 %). Mice were fed *ad libitum* with a chow diet, except for the Zileuton experiment, described below. Cage bottoms were covered with autoclaved bedding and enrichment material.

All mice were sacrificed by slow rising concentration of CO_2_ followed by confirmation of death by cervical dislocation. Tools used to obtain samples were sanitized using 70% ethanol before and after handling.

### Primary cells

For confocal microscopy experiments, pMФ were obtained as described previously^12^ via peritoneal lavage (see below). The cells were either processed for flow-cytometric cell sorting or plated in complete medium (RPMI 1640 supplemented with 10% FCS, 100 U.ml^-1^ penicillin, 10 µg.ml^-1^ streptomycin and 400 µM L-glutamine (Invitrogen)) for *in vitro* culture. Cells were plated and left to adhere for 3 h at 37 °C, 5% CO_2_, then washed two times with warm complete medium.

### Cell lines

HEK293T cells and RAW264.7 cells were cultured in ‘D10’ (DMEM (Invitrogen) containing 10 % FCS (Gibco Life Technologies, Dublin, Ireland), 100 U.ml^-1^ penicillin, 10 µg.ml^-1^ streptomycin at 37 °C and 5 % CO2. Cells were passaged every two days, when 80-90 % confluence was reached.

### Peritoneal lavages

Peritoneal cells were obtained via peritoneal lavage as described previously^12^ using ice-cold 5 ml lavage solution (PBS (Invitrogen) supplemented with 5 mM EDTA and 4% foetal calf serum (FCS)). Cells were counted in triplicate using Muse cell analyser (Luminex Corporation). For lipid analysis experiments, FCS was omitted from the lavage buffer.

For collection of cell free lavage fluid, mice were lavaged as above with 1 mL of ice-cold PBS (calcium and magnesium free), without FCS or EDTA. The collected cell suspension was centrifuged for 5 minutes at 350xg at 4°C and liquid was removed carefully so as not to disturb the cell pellet. Collected liquid was frozen and stored at - 80°C for protein analysis or lipid extraction.

### Mass spectroscopy

For all lipid analysis, cells were sorted to high purity (>98%) on a FACSAria III (BD biosciences, Berkshire, UK) as described later.

### Lipids extraction for analysis of oxylipins

Oxylipin extraction was performed as previously described^15^. Cells or cell free lavages were defrosted on ice and transferred to glass universals. Cell pellets were resuspended in 1 mL of PBS and cell free lavage samples were measured for future analysis and made up to 1mL with HPLC grade water. Internal standards were added to each sample at concentrations indicated in **Supplementary Table 4**. 2.5 mL of extraction solution (1 M acetic acid/2-isopropanol/hexane (2:20:30, v/v/v)) was added per 1 mL of sample. Samples were vortexed for 1min, followed by addition of 2.5 mL of hexane and further vortexing for 1 min. Samples were also mixed by inverting the tube before they were centrifuged for 5 min at 1500 rpm at 4°C. Lipids were recovered in the upper organic solvent layer which was collected into separate glass universal using a glass pipette. 2.5 mL of hexane was added to the remaining bottom layer and the sample was vortexed for 1min and then centrifuged for 5min as indicated above. The top layer was collected and combined with the previous collection and dried in RapidVap drier at vacuum 200, speed 30 and at 30 °C. The remaining aqueous layer was re-extracted using Bligh & Dyer method^16^. Specifically, 2.5 mL methanol and 1.25 mL of chloroform was added to the samples, followed by vortexing for 1min and a further addition of 1.25 mL of chloroform and 1.25 mL of water with vortexing steps in between. Samples were spun as indicated above. The bottom chloroform layer was collected and combined with the upper layers from the hexane/IPA extraction. Samples were resuspended in 100 μL methanol for LC-MS/MS analysis.

### LC-MS/MS targeted analysis of oxylipins

Targeted analysis of oxylipins was performed on a Nexera liquid chromatography system (Nexera Xa, Shimadzu coupled to a 6500 Qtrap mass spectrometer (AB Sciex). Liquid chromatography was performed at 45 °C using a Zorbax Eclipse Plus C18 (Agilent Technologies) reversed-phase column (150 × 2.1 mm, 1.8 μm) at a flow rate of 0.5 ml/min over 22.5 min. Mobile phase A was (95% HPLC water/5% mobile phase B; v/v and 0.1% acetic acid) and mobile phase B was acetonitrile/methanol (800 ml + 150 ml; and 0.1% acetic acid). The following linear gradient for mobile phase B was applied: 30% for 1 min, 30–35% from 1 to 4 min, 35–67.5% from 4 to 12.5 min, 67.5–100% from 12.5 to 17.5 min and held at 100% for 3.5 min, followed by 1.5 min at the initial condition for column re-equilibration. The injection volume was 5 µL. Lipids were analyzed in scheduled multiple reaction monitoring mode with a detection window of 55 s for each analyte. Ionization was performed using electrospray ionization in the negative ion mode with the following MS parameters: temperature 475 °C, N_2_ gas, GS1 60 psi, GS2 60 psi, curtain gas 35 psi, ESI voltage −4.5 kV. Cycle time was 0.4 s.

**Supplementary Table 4** contains the list of abbreviations and formal names for lipids investigated in our panel. It also contains a full list of the internal standards and oxylipins along with calibration range, MRM transitions and corresponding MS parameters (collision energy, declustering potential).

### LC-MS/MS targeted analysis of CysLTs

Targeted analysis of CysLTs was performed on a Nexera liquid chromatography system (Nexera Xa, Shimadzu coupled to a 6500 Qtrap mass spectrometer (AB Sciex). Liquid chromatography was performed at 30 °C using a BEH C18 (Waters) reversed-phase column (50 × 2.1 mm, 1.7 μm) at a flow rate of 0.6 ml/min over 15 min. Mobile phase A was water + 0.1% formic acid and mobile phase B was acetonitrile + 0.1% formic acid). The following linear gradient for mobile phase B was applied: 5% for 1 min, 5–53% from 1 to 9.5 min, 53-76% from 9.5 to 11 min, 76–100% from 11.1 to 11.1 min and held at 100% until 12.1 min, followed by 3 min at the initial condition for column re-equilibration. The injection volume was 10 µL. Lipids were analyzed in scheduled multiple reaction monitoring mode. Ionization was performed using electrospray ionization in the negative ion mode with the following MS parameters: temperature 600 °C, N_2_ gas, GS1 20 psi, GS2 5 psi, curtain gas 35 psi, ESI voltage −4.5 kV.

For both oxylipin and CysLT analysis, peak areas for lipids were integrated and normalized for deuterated internal standards and quantification was performed using external calibration curves with authentic standards for all investigated oxylipins. Data were acquired using Analyst (V1.6). Integration and quantification was performed using MultiQuant software (version 3.0.2, AB Sciex Framingham, MA, USA). All chromatograms from tissue/cell samples were manually checked for peak quality and excluded from analysis if they fell below the limit of 5:1 (signal:noise, LOQ) with a minimum of six points across each peak. Where lipids fell below LOQ, a zero value was recorded, but replaced with 50%LOQ for statistical analysis.

### Untargeted lipidomic analysis

Global lipidomic analysis was carried out on an Accela liquid chromatography system coupled to an Orbitrap Elite mass spectrometer (Thermo Fisher Scientific). Liquid chromatographic separation was performed at 30°C using an Accucore C18 (Thermo Fisher Scientific) reversed phase column (150 × 2.1 mm, 2.7 μm) with solvent gradient of mobile phase A (water/acetonitrile; 80/20; v/v; 4 mM ammonium acetate and 0.1 % glacial acetic acid) and B (acetonitrile/isopropanol; 30/70; v/v; 4 mM ammonium acetate and 0.1 % glacial acetic acid) at 0.4 mL/min over 60 min. The linear gradient of B was 16 - 60% for 12 min, 60 - 72% from 12 min - 19 min, 72 - 84% from 19 min - 42 min, 84-98% from 42 min - 51 min and held at 98% for another 6.5 min before re-equilibration. MS conditions were as follows for analysis in positive ESI ionization mode: resolution 60,000 at 400 *m/z* (providing approx. 3 scans/sec) HESI-II temperature 400°C, N_2_ as drying gas, sheath gas flow 37 arbitrary units (au), auxiliary gas flow 15 au, sweep gas flow 1 au, capillary temperature 320°C, spray voltage + 4.0 kV, S-lens RF level 62 %. Lock mass was *m/z* 391.2843. For analysis in negative ESI ionization mode: resolution 60,000 at 400 *m/z*, HESI-II temperature 350°C, N_2_ as drying gas, sheath gas flow 37 au, auxiliary gas flow 15 au, sweep gas flow 2 au, capillary temperature 320 °C, spray voltage – 3.5 kV, S-lens RF level 69 %. Lock mass was *m/z* 265.1479.

### XCMS processing

The .RAW files produced from the LC-MS runs were converted to .mzML format applying ProteoWizard 2.1^17^ followed by deconvolution and peak alignment applying XCMS using a previously described method ^18^. Results were then queried against two databases, Human Metabolome Database (HMDB) and LIPID MAPS using the database search program MS search with mass tolerance of 5 ppm. As only precursor *m/z* values were used, putative names were not assigned, instead, lipids were assigned putatively to categories that match the most common assignments for any individual *m/z* value. This approach allows behaviour of individual lipid categories to be assessed as an initial step.

### Targeted sphingolipid analysis

Sphingolipid analysis was performed using LC-MS/MS on a Shimadzu LC-30 AD binary pump system with a Shimadzu SIL-30 AC autosampler, coupled to a 6500 QTrap (AB SCIEX™, Warrington, UK) operating in triple quadrupole mode. Lipid extracts were separated on an Accucore C18 column (150 x 2.1 mm, 2.6 µm) using an isocratic mobile phase consisting of Phase A (H₂O, 80:20 v/v) and Phase B (IPA, 70:30 v/v) with 4 mM ammonium acetate, at a flow rate of 0.4 mL/min over 35 minutes. The column temperature was maintained at 25 °C, and samples were kept at 4 °C.

Lipid standards were used to establish multiple reaction monitoring (MRM) conditions and to generate calibration curves for quantification. After optimisation of instrument parameters for each sphingolipid class, standard curves were prepared by serial dilution of the internal standards and analysed under the final LC–MS/MS method. Because sphingolipid species show comparable ionisation efficiencies across different acyl-chain lengths, a C12-based standard was used for quantification of all species, as previously reported^19^.

The internal standard mixture (Cer/Sphingoid Internal Standard Mixture II; Avanti Polar Lipids, Alabaster, AL, USA) and its composition, a full list of analytes, internal standards, and their DP, CE, and MRM transitions are provided in **Supplementary Table 4**.

The limit of quantification (LOQ) was defined as a signal-to-noise ratio ≥5 with at least six data points per chromatographic peak. Limits of detection (LOD) and LOQ were determined from serial dilutions of the internal standards. Lipids with concentrations below the detection limit (recorded as zero) were imputed at 50% of the LOD.

### Antibody and drug treatments

Total of 20 μg of Ultra-LEAF purified anti-mouse/human IL-5 antibody (TRFK-5 clone, Biolegend) was injected intraperitoneally to mice. Control mice were injected with 20 ug of Ultra-LEAF purified Rat IgG1 isotype-matched control antibody (Biolegend, #400432) in total of 150 ul of sterile, pyrogen, Mg+ and Ca+ free PBS. Lavages were collected 72 hours after injection.

NGB-2 NutraGel (AIN-93, Bacon Flavor (Complete Nutrition), LBS Biotechnology) was liquidised in warm water bath. Total of 10 mg of Zileuton (A-64077, Stratech Scientific Ltd) powder was mixed into 2 oz of liquidized sterile NGB-2 NutraGel and fed to up to 6 mice when returned to solid state. Treatment was repeated every 24 hours. Mice had no access to any other food source during the experiment. The amount of NutraGel was calculated to deliver minimum of 70 mg/kg of Zileuton per mouse, based on an average mouse body weight of 22 g and daily intact of about 9 g of NutraGel. NutraGel without any addition was used as a control. NutraGel handling in the laboratory was performed in class II Biological Cabinet.

Indomethacin was initially dissolved in ethanol and diluted 2000-fold for final dose of 1 mg/kg concentration in 0.2 ml of sterile, pyrogen, Mg+ and Ca+ free PBS. PBS containing 0.05 % Ethanol was given in parallel to control animals. Delivered via oral gavage, twice, with an interval of 24 hours in total volume of 200 ul. Peritoneal lavage was collected 24 hours after second gavage.

### Preparation of lentiviral vectors and infectious lentiviral particles

Lentiviral vectors were generated as previously described ^20, 21^. In brief, shRNA against target genes were inserted using ‘In-fusion’ cloning kit (Clontech) in EGFP expressing vector, pHR’SIN-cPPT-SEW^6^ and the lentivirus particles were produced using HEK293T cells and purified using a sucrose gradient ultracentrifugation step. Lentiviral particles were resuspended in Aim-V medium (Life Sciences) and stored at -80 °C until further use. The sequence of the primers used to generate the shRNA: *Smpd1* Forward: 5’-AGA AAA GCC TTG TTT GGA GCC TCC CAG ATG CTA ATA CTC GAG TAT TAG CAT CTG GGA GGC TCC TTT TTG TTT AAA CGA ATT CCT GCA-3’, Smpd1 Reverse: 5’-TGC AGG AAT TCG TTT AAA CAA AAA GGA GCC TCC CAG ATG CTA ATA CTC GAG TAT TAG CAT CTG GGA GGC TCC AAA CAA GGC TTT TCT-3’; *Gba2* Forward: 5’-AGA AAA GCC TTG TTT GTA CAA ATC TGC ACT GTT CAA CTC GAG TTG AAC AGT GCA GAT TTG TAC TTT TTG TTT AAA CGA ATT CCT GCA-3’, *Gba2* Reverse: 5’-TGC AGG AAT TCG TTT AAA CAA AAA GTA CAA ATC TGC ACT GTT CAA CTC GAG TTG AAC AGT GCA GAT TTG TAC AAA CAA GGC TTT TCT-3’.

### In vitro RAW264.7 cells lentiviral infection

RAW 264.7 cells were plated into a 12-well plate (4×10^5^ cells in 400 µl D10 complete medium) and left to adhere for 4 h before addition of 100 µl of lentivirus. Additional D10 medium was added after 24 h. Five days after infection, the GFP^+^ RAW264.7 cells were purified using flow-cytometry sorting on FACSAria III (BD). 2×10^6^ cells per sample were used for downstream targeted sphingolipid analysis.

### Flow-cytometry, Imaging flow-cytometry and flow cytometric cell sorting

Flow-cytometric staining was conducted using standard protocols^6, 22–25^. In brief, the genotype of the individual peritoneal Gata6-WT and KO^mye^ lavage samples was confirmed phenotypically using F4/80-APC (Biolegend) and either CD73-ef450 (eBioscience) or TIM-4 (BD Biosciences). Flow cytometry was performed on Cyan (Beckman Coulter) or Attune (Thermofisher) flow cytometers and analysed with FlowJo software. Imaging flow cytometry was performed on Amnis ImageStream XMkII (Luminex) instrument and analysed with IDEAS (v6.2) software.

For flow-cytometric cell sorting, lavages from identically genotyped mice were pooled and stained with F4/80-APC, CD73-ef450 and MHCII-PercpCy5.5 (BD biosciences) for sphingolipid analysis or with F4/80-Pacific Blue, TIM-4-PE and Siglec-F-AF647 for eicosanoid experiments. Cells were sorted on a FACSAria III (BD biosciences, Berkshire, UK) for single events (via pulse width or FSC_height_ vs FSC_area_ respectively) with the defined flow-cytometric phenotype: pMФ Gata6-WT: F4/80^high^, MHCII^low^, CD73^high^; pMФ Gata6-KO^mye^ - F4/80^+^, MHCII^low^, CD73^+^ or pMФ Gata6-WT: F4/80^high^, TIM-4^+^; pMФ Gata6-KO^mye^ - F4/80^+^, TIM-4^+^, eosinophils: SSC^high^, Siglec-F^+^, mast cells SSC^high^ , Siglec-F^-^, F4/80^-^, TIM-4^-^. Cell purities were verified by flow cytometry before the processing the samples.

### Immunofluorescent microscopy

Cells were plated in complete medium on cover glass (VWR, thickness no. 1) and left to adhere for 3 h before washing 3 times with complete RPMI medium. Following indicated treatment, cells were fixed for 15 min with 4% PFA and permeabilized and blocked for 30 min with PBS containing 0.1% (v/v) Triton X-100 (Sigma), 10% (v/v) FCS and 10% (v/v) rabbit serum. The cells were stained overnight with F4/80-AF488 (clone BM8, Biolegend). BODIPY staining was performed accordingly to the manufacturer’s instruction (BODIPY 493/503, Thermofisher). DAPI (Thermofisher) was applied as counterstain and cells were mounted on microscope slides (Thermoscientific) using fluorescent mounting medium (Dako). Pictures were taken using LSM800 confocal laser scanning microscope (Zeiss).

### Quantitative real-time PCR

Total RNA was isolated from cells using RNeasy Mini or Micro Kit (Qiagen) following the manufacturer’s instructions and at least 350 ng were reverse transcribed using the High-Capacity cDNA Reverse Transcription kit (Applied Biosystems). mRNA levels were quantified using a ViiA™ 7 Real-Time PCR System (Applied Biosystem) and Power SYBR Green PCR Master Mix (Thermofisher). The gene expression values were normalized to *Ywhaz* expression and normalized to WT control.

### Protein level measurements

Composition of CCL11, CCL24, IL-5, IL-33 in the peritoneal cell free lavages was measured using ELISAs (Mouse DuoSet CCL11, CCL24 and IL-33, all from R&D Systems; Mouse IL-5 ELISA MAX Deluxe, Biolegend) accordingly to manufacturer protocols. The relative levels of 40 cytokines in cell free lavages was measured using Proteome Profiler Array - Mouse Cytokine Array Panel A (R&D Systems) accordingly to manufacturer protocols.

### Network and pathway analysis

The microarray data used in this study were previously published^6^. To investigate the biological relationships and functional significance of the differentially expressed genes, network and pathway analyses were conducted using QIAGEN’s Ingenuity Pathway Analysis (IPA®, QIAGEN, Redwood City, CA, USA, www.qiagen.com/ingenuity).

### Data analysis

Statistical data analysis was performed using Prism 10.4.0 (GraphPad Software) or R version 3.6.2^84^ (Sphingolipids data). Parametric tests were employed for normally distributed data, while non-parametric tests were utilized for skewed distributions. The exact statistical methods used for the experiments are indicated in the figures, figure legends and methods section.

For the untargeted lipid analysis, XCMS-analysed data were processed using both a univariate and multivariate approach. Significantly different lipids were identified using an unpaired parametric t-test (Student t-test, unequal variances). *P*-values were adjusted for multiple comparisons, using the Benjamini-Hochberg method. Differential expression was classified as significant when P < 0.1 and |FC| > 1.5. These thresholds were chosen to avoid missing potentially meaningful differences due to the data complexity and limited replicate number, knowing that targeted analysis would follow to validate this initial conclusions^26^. Unsupervised PCA and supervised OPLS methods were used to obtain group clusters using the program Simca (version 14.0.1.0, Umetrics AB, Umeå, Sweden, 2013).

For statistical analysis of targeted sphingolipid data, each lipid species was tested individually for differences between the two groups (Gata6-WT vs Gata6-KO^mye^) using an unpaired two-tailed Student’s t-test. P-values were adjusted for multiple comparisons using the Benjamini–Hochberg method, and adjusted P < 0.05 was considered significant. All statistics were performed using GraphPad Prism 6 software.

## Results

### *Gata6*-deficiency leads to sphingolipid accumulation in resident peritoneal macrophages

To evaluate the impact of tissue resident MФ programming on lipid metabolism, we focused on resident pMФ (**Figure S1A**) and the role of GATA6, using mice with a myeloid-restricted *Gata6*-deficiency (*Gata6*-KO^mye^)^6^. First, BODIPY staining of pMФ, revealed a significant increase in neutral lipid content in *Gata6*-KO^mye^ when compared to *Gata6*-WT pMФ, which appeared to be mostly located in lipid-rich vesicles, and was evident both in freshly isolated cells and following 12 hours in culture (**Figure 1A-C**). Having established the presence of lipid-rich vesicles in the *Gata6*-deficient cells, we undertook a preliminary Differential Interference Contrast (DIC) and coherent anti-Stokes Raman scattering (CARS) microscopy analysis of the representative pMФ, from each genotype. This small-scale assessment confirmed the characteristic morphology and presence of the vesicles in *Gata6*-KO^mye^ pMФ (**Figure 1D**, red vesicles, and **S1B**).

**Figure 1:**
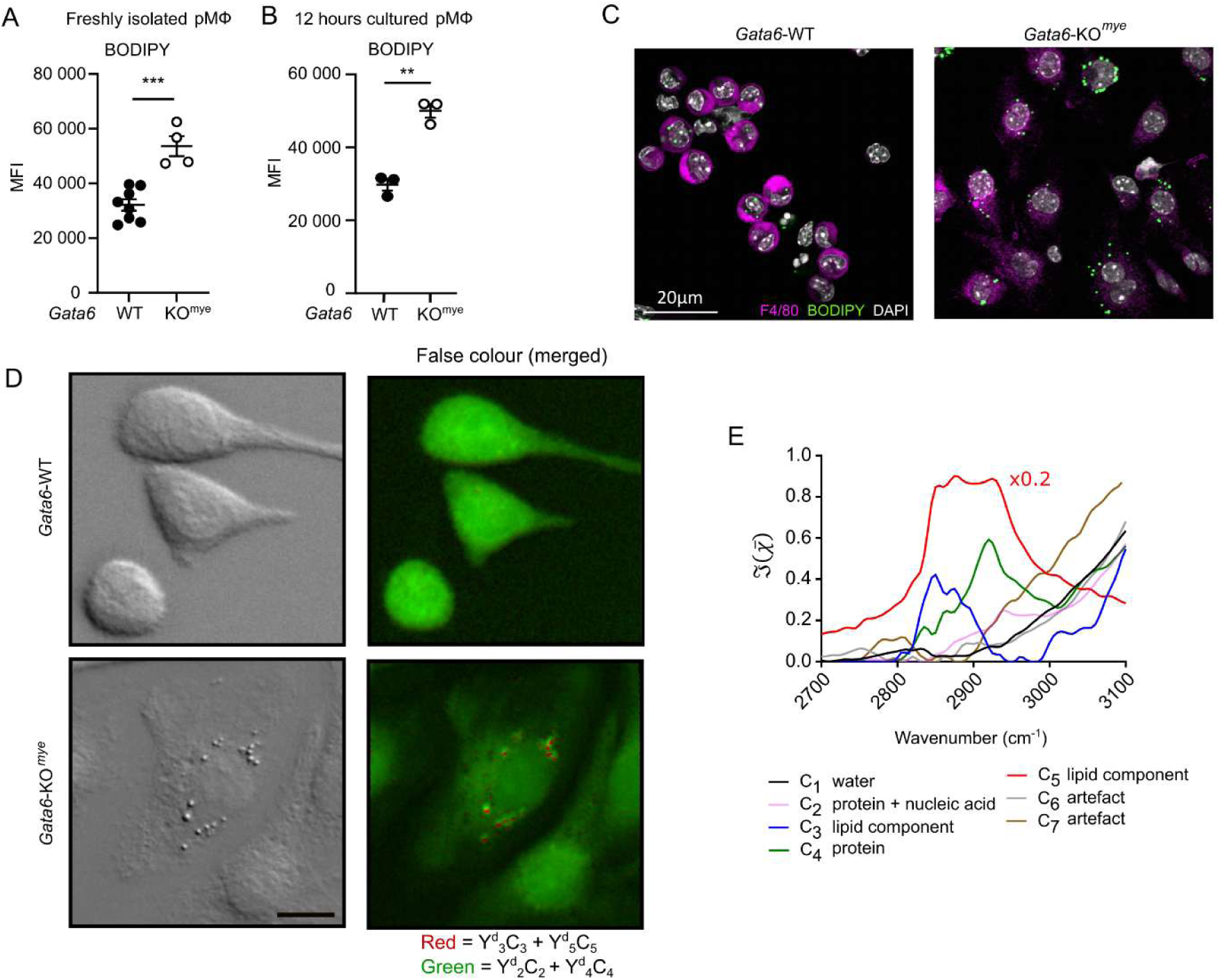
GATA6 controls lipid accumulation in resident peritoneal macrophages. Mean fluorescence intensity (MFI) flow-cytometric analysis of BODIPY staining performed on freshly isolated **(A)** and 12 hour cultured **(B)** *Gata6*-WT and *Gata6*-KO^mye^ pMФ from male mice. Each symbol represents an individual mouse. Statistical significance was determined using unpaired t-test, **p<0.01, ***p<0.001. **(C)** Confocal microscopy analysis of BODIPY staining localisation in *Gata6*-WT and *Gata6*-KO^mye^ pMФ. **(D)** DIC images (left) from *Gata6*-WT and *Gata6*-KO^mye^ pMФ and corresponding false colour images generated from the overlay of FSC^3^ concentration maps of components C_3_ (blue, lipid component), C_4_ (green, protein), C_5_ (red, lipid component). Both images are on the same RGB scale, from 0 to a colour maximum scale M (vol/vol) with M_red_ = 0.167, M_green_ = 0.082. Scale bar: 10 μm. Analysis was performed on fields of view highlighted in black boxes in Figure S1B. **(E)** Susceptibility spectra from the FSC^3^ analysis. c_5_ has been multiplied by 0.2 to allow all components to be seen clearly on a single graph.

To identify the chemical composition of these vesicles, hyperspectral CARS image stacks were obtained and analysed using in-house developed Hyperspectral Image Analysis (HIA) software^27–31^. The data was factorised into 7 components with associated CARS susceptibility spectra and spatially resolved concentrations using FSC^3,27–31^. Based on the retrieved spectra (**Figure 1E**) and the spatial profile of the concentration maps (**Figure S1C**), two lipid components were identified, labelled as c_3_, c_5_ in their spectral profile having concentration maps C_3_, C_5_. c_3_ shows a narrower profile with a stronger peak around 2,850 cm^-1^ wavenumber (characteristic of the CH_2_ symmetric stretch vibration) resembling the spectrum of more ordered, saturated lipids. c_5_ on the other hand, has a broader profile with a pronounced shoulder around 2,930 cm^-1^ (due to a combination of CH_3_ stretch and CH_2_ asymmetric stretch vibrations) indicative of disordered, unsaturated lipids^31, 32^. The remaining components were nominally attributed as c_1_: water, c_2_: protein + nucleic acids, c_4_: protein. The spectral shape of c_6_ and c_7_ and their concentration maps C_6_ and C_7_, indicated that these were not physically meaningful chemical components and were attributed to artefacts. The component spectra exhibit a rising tail above 3000 cm^-1^, indicative of water as shown by component c_1_. This is indicative of water being present across the imaged volume in each voxel. Therefore, prior to generating false colour images, the concentrations C_2_ to C_7_ were scaled to represent dry values, by removing the water contribution, as described in our recent work^29, 33^. Briefly, for each component, we calculated the dry volume fraction γ*^d^_i_*, to obtain the corresponding dry vol/vol concentration γ*^d^_i_C_i_*. Details on the determination of γ*^d^_i_* are given in the supplement (**Supplementary Methods**). False colour dry concentration maps of the FSC^3^ components containing lipids γ*^d^_3_C_3_* + γ*^d^_5_C_5_* (red) and proteins γ*^d^_2_C_2_* + γ*^d^_4_C_4_* (green) were created for comparison and correlation with DIC images (**Figure 1D**). These concentration maps indicate a higher amount of lipid present in the cytoplasm of *Gata6*-KO^mye^ pMФ as compared to *Gata6*-WT. Furthermore, the spatial pattern of the lipid components correlates with the round-shaped vesicles seen in the DIC images of *Gata6*-KO^mye^, identifying these as lipid-rich vesicles (**Figure 1E and S1C**). Together, results obtained from BODIPY staining and CARS microscopy indicate accumulation of lipids in vesicles in *Gata6*-KO^mye^ pMФ.

Next, to identify which lipids had accumulated in these cells, an untargeted high resolution liquid chromatography-mass spectrometry (LC/MS) screen was performed using total lipid extracts obtained from freshly isolated *Gata6*-WT and *Gata6*-KO^mye^ pMФ (**Supplementary Table 1**). Putative annotation of signals to lipid categories suggested that the lipidome of *Gata6*-KO^mye^ pMФ was distinct from WT, with significant upregulations in lipid levels across several different categories (**Figure S2A**). Sphingolipids (SL) and glycerophospholipids had the highest proportion of lipids changed, primarily upregulated, in *Gata6*-KO^mye^ pMФ, at 15.2% changed (14.1% upregulated, 0.9% downregulated and 0.2% unique to *Gata6*-KO^mye^ cells) and 10.5% changed (9.6% upregulated, 0.8% downregulated, and 0.1% unique to *Gata6*-WT cells), respectively (**Figure S2**). Mirror plots show the changed ions between the two genotypes and emphasize the predominant upregulation seen in the *Gata6*-KO^mye^ cell (**Figure S2B**) and loading plots show the significantly altered lipids by category (**Figure S2C**). SL are divided into main classes including, ceramides (Cer), lactosylceramide (LacCer), sphingomyelin (SM), and metabolites such as ceramide-1-phosphate (C1P). Thus, to characterise the specific impact on SL modulation in *Gata6*-KO^mye^ pMФ, we next performed a targeted LC-MS/MS analysis of SL molecular species in *Gata6*-WT and *Gata6*-KO^mye^ pMФ. As for our untargeted analysis, this demonstrated that most SL analysed across several classes were significantly increased in *Gata6*-KO^mye^ pMФ, and this was regardless of fatty acyl chain length or saturation (**Figure 2A, B, Figure S3A**).

**Figure 2:**
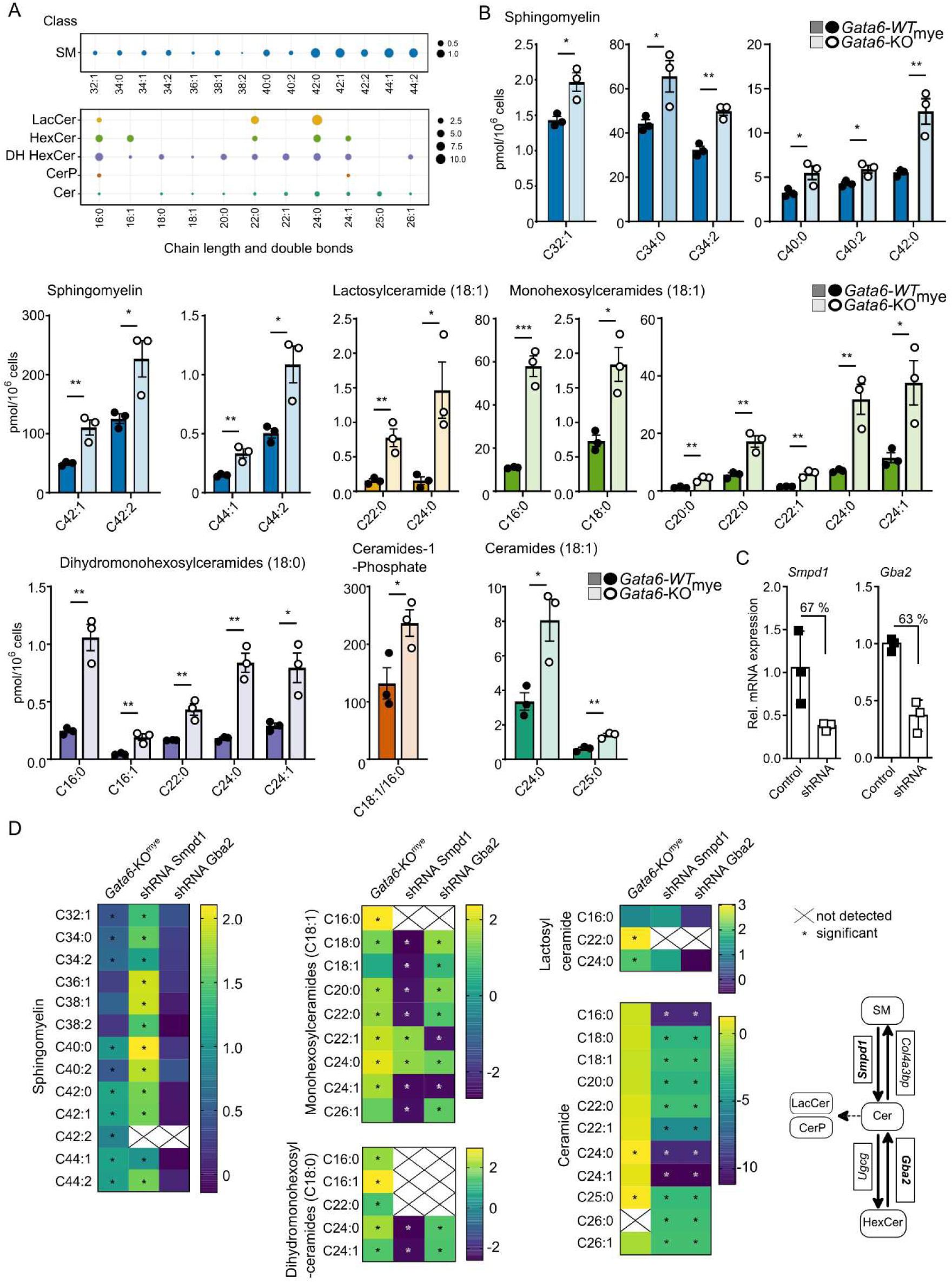
Dysregulation of sphingolipids in *Gata6*-deficient resident pMФ. **(A)** Log_2_FC (*Gata6*-KO^mye^ / -WT pMФ) of sphingolipids by lipid class and sub-class by chain-length. Log_2_FC is indicated by spot size. **(B)** Bar chart representation of sphingolipids by lipid class and sub-class by chain-length significantly changed in *Gata6*-KO^mye^ pMФ compared to *Gata6*-WT. Statistical significance determined using t-test. Each lipid was analysed individually, without assuming a consistent SD, the graphs depict the mean±SEM, each symbol represents an individual mouse. Colours correspond to specific lipid category, with *Gata6*-KO^mye^ in paler shading. **(C)** qPCR analysis of *Smpd1* and *Gba2* expression in RAW264.7 cells following infection with lentivirus containing shRNA specifically directed against the target genes or non-silencing control shRNA (Control). Analyses were performed on cells after selection by flow-cytometric sorting to high purity (>95%) on reporter gene expression (eGFP^+^), 7 days after infection. Data expressed as mean±SEM from three independent experiments. **(D)** Heatmaps of significantly changed sphingolipids in RAW264.7 cells (log2 FC) following infection with lentivirus containing shRNA against Smpd1 or Gba2 and changes to that lipid in *Gata6*-KO^mye^ pMФ. The heatmaps show the difference (fold-change) between the indicated group and their respective controls: nd - levels of lipid not exceeding background, ns – not significant from its respective control. *p<0.05, **p<0.01, ***p<0.001.

### Sphingolipids in pMФ are regulated by discrete transcriptional changes present in *Gata6*-deficient cells

To define how the loss of GATA6 alters SL levels, we next interrogated transcriptomic data from *Gata6*-WT and *Gata6*-KO^mye^ pMФ^6^. Here, causal Ingenuity Pathway Analysis Network Analysis predicted several lipid metabolism gene expression dysregulations in *Gata6*-KO^mye^ pMФ. In particular, the “quantity of sphingolipid”, the “quantity of glycosphingolipid”, and the “concentration of sphingomyelin” were all predicted to be activated (**Figure S3B**), indicating that transcriptional regulation directly contributed to SL accumulation in these cells. Analysis of the genes involved in the SL metabolic pathway showed significant changes to expression of multiple key genes (**Figure S3C,D**) including serine palmitoyltransferase long chain base subunit 2 (*Sptlc2*), a key enzyme in sphingolipid biosynthesis^34^. In contrast, sphingomyelin phosphodiesterase 1 (*Smpd1*), which catalyses conversion of SM to Cer^35, 36^, was significantly downregulated in GATA6-deficient pMФ (**Figure S3D**), which could explain the accumulation of SM in these cells. Similarly, conversion of Cer to HexCer is driven by UDP-glucose ceramide glycosyltransferase (*Ugcg*) and reversed by glucosylceramidase beta 2 (*Gba2*)^37, 38^. While regulation of these lipid species is very complex, expression of both of these genes was significantly increased and decreased, respectively, consistent with the observed HexCer accumulation in *Gata6*-KO^mye^ pMФ (**Figure S3C**). To discriminate between glucosylceramide (GlcCer) and galactosylceramide (GalCer), LC-MS/MS conditions were modified as described by Shaner et al.^19^; GalCer was not detected in pMФ from either genotype, confirming that the HexCer signal represents GlcCer (**Supplementary Methods, appendices 4 and 5**).

Given the significant downregulation of *Gba2* and *Smpd1* in *Gata6*-KO^mye^ cells, and the observed lipid changes, such as concurrent changes in SL, we hypothesized that these changes in *Gata6*-KO^mye^ cells may be causative. To confirm the impact of *Gba2* and *Smpd1* downregulation on SL balance in macrophages, we next employed the macrophage cell line, RAW264.7, which exhibited similar expression of both genes to WT pMФ (**Figure S3E**). RAW264.7 cells were transduced with lentivirus containing shRNA^20, 21^ directed against *Gba2* and *Smpd1*. This led to a significant downregulation of the target genes, by 63% and 67%, respectively, as compared to the non-silencing control (**Figure 2C**). The reduction of *Gba2* and *Smpd1* expression seen mirrored the fold-change difference of these genes observed between *Gata6*-WT and -KO^mye^ pMФ (*Gba2* 52% and *Smpd1* 51%). Targeted LC-MS/MS analysis demonstrated that downregulation of *Smpd1* alone was sufficient to induce significant LacCer, SM and DH SM increases in RAW264.7 cells similar to changes observed in *Gata6*-KO^mye^ pMФ (**Figure 2D**). In contrast, the levels of Cer and HexCer were decreased in *Smpd1* shRNA treated RAW264.7 cells in line with the role of *Smpd1* in metabolising ceramides from SM. *Gba2* downregulation also affected SL content in RAW264.7 cells, but mostly only HexCer and DH HexCer classes (**Figure 2D**). Data is shown in **Supplementary Tables 2 and 3**. Thus, the specialisation of pMФ following the expression of *Gata6* seems to be a major checkpoint allowing for the correct expression of SL metabolism genes, in particular *Smpd1* and *Gba2.* Together, these data suggest that tissue specialisation of pMФ controls SL metabolism, driven by transcriptional changes that occur during this specialisation process.

### The absence of peritoneal macrophage specialisation alters oxylipin biosynthesis

Next, we examined for functional implications of the altered SL metabolism that was observed in pMФ lacking GATA6. A previous study found that sphingolipid and oxylipin biosynthetic enzymes are functionally linked in peritoneal-derived mast cells^39^. Specifically, upregulation of endoplasmic reticulum membrane protein ORM1-like protein 3 (ORMDL3) and serine palmitoyltransferase1 (SPT1) were shown to be functionally linked with increased generation of leukotrienes (LT), with the two proteins physically associating with 5-LOX to drive LT synthesis^39^. Furthermore, Cer, C1P and LacCer can activate cytosolic phospholipase-A2α(PLA_2_)^40–42^, an enzyme that releases oxylipin precursor fatty acyls (FA), while D-erythro-analogs of sphingosine have the opposite effect on PLA_2_^43^. Oxylipins, such as prostaglandins (PG) and cysteinyl leukotrienes (CysLTs), are bioactive lipid mediators, which play immunomodulatory functions in homeostasis and inflammation^44^. We previously demonstrated altered PG levels in cultured *Gata6*-KO^mye^ pMФ, which significantly alter their immune responses to LPS^12^. Extending this, targeted LC-MS/MS analysis of 93 oxylipins was performed using freshly isolated *Gata6*-WT and -KO^mye^ pMФ at homeostasis (**Figure 3A, and Figure S4A, Supplementary Table 2**). A total of 16 were detected and while most were unchanged, increased thromboxaneB_2_ (TxB_2_) and reduced 6-ketoPGF1α was seen in *Gata6*-KO^mye^ pMФ, as previously reported^12^ (**Figure 3A**). Peritoneal eosinophils and mast cells were isolated from the same samples and found to show no significantly altered oxylipins between *Gata6*-WT and -KO^mye^ (**Figure S4A**). *Gata6* is not expressed in these cell types, providing a potential explanation.

**Figure 3:**
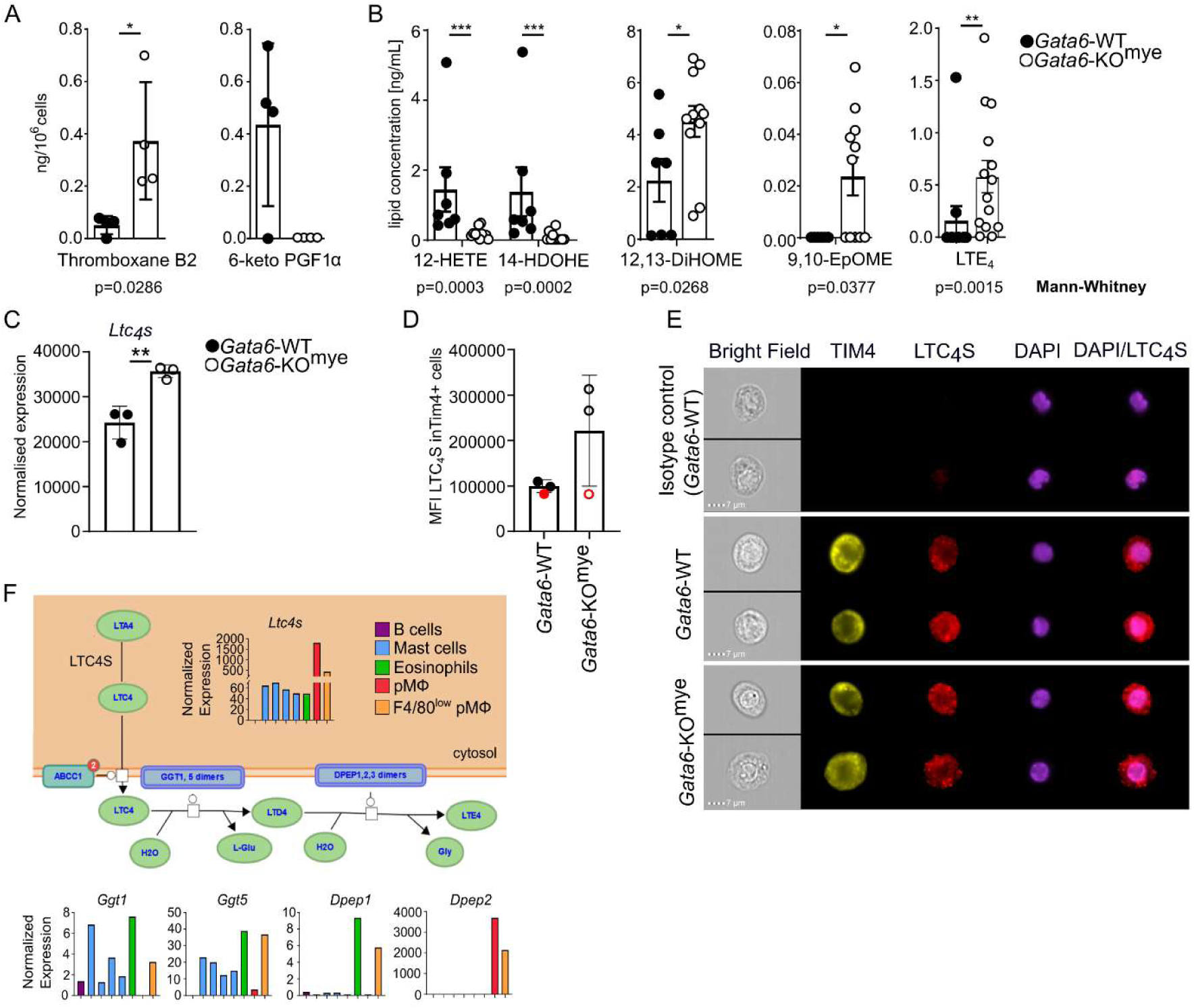
Differential eicosanoid production in the absence of myeloid GATA6. **(A)** Bar graphs showing expression of changed eicosanoids inside freshly isolated *Gata6*-WT and - KO^mye^ pMФ and **(B)** in cell free peritoneum, measured using LC-MS/MS. Statistical significance was determined using Mann-Whitney test. **(C)** Relative expression of *Ltc4s* in *Gata6*-WT and *Gata6*-KO^mye^ pMФ, measured by previously published microarray. **(D)** ImageStream analysis of LTC_4_S in *Gata6*-WT and -KO^mye^ pMФ from 2 independent experiments indicated by the symbol colour and **(E)** a representative gallery from one experiment. **(F)** Immgen gene expression of enzymes involved in LTC_4_ to LTE_4_ metabolism in different cell types in the peritoneal cavity. *P*-value, *<0.05, **<0.01, ***<0.001. Data in bar graphs represents mean±SD, and each symbol represents an individual mouse, apart from (A) where multiple mice (10-18) of the same genotype were pooled for each point.

However, oxylipins are primarily secreted from cells and detected in extracellular media in larger amounts than in isolated cells. Therefore, cell free peritoneal lavage from *Gata6*-WT and -KO^mye^ mice were next analysed. In total, 15 oxylipins were detected, with the 12/15-LOX metabolites 12-HETE and 14-HDOHE being significantly reduced (**Figure 3B, S4B**), consistent with the reduced expression of *Alox15* in *Gata6*-KO^mye^ pMФ (**Figure S4D,E**). Three oxylipins were significantly increased in *Gata6*-KO^mye^ lavage, namely 12,13-DiHOME, 9,10-EpOME and LTE_4_. (**Figure 3B**). Example chromatograms for LTE_4_ are provided in **Figure S4C**. Significant increases in *Tbxas1* and *Ptgis* gene expression were seen in *Gata6*-KO^mye^ pMФ, consistent with elevated TxB_2_ and 6-ketoPGF1α (**Figure S4D,E**). Expression of LTC_4_ synthase (*Ltc_4_s)*, which conjugates LTA_4_ to reduced glutathione, forming LTC_4_, which is then further metabolised to LTE_4_, was significantly elevated in *Gata6*-KO^mye^ pMФ (**Figure 3C**). LTC_4_S was also detected at the protein level using imaging flow cytometry (**Figure 3D,E**). Immgen Skyline^45^ analysis of peritoneal immune cells gene expression showed that classic pMФ, defined by F4/80^+^ICAM^+^CD5^-^CD19^-^CD43^-^ are the primary cell type expressing LTC_4_S in the peritoneal cavity (**Figure 3F**). Further biosynthesis of CysLTs may also be mediated by small peritoneal macrophages (F4/80^low^pMФ), eosinophils and mast cells, which demonstrate the highest expression of gamma-glutamyltransferase (GGT) and dipeptidase (DPEP), membrane bound enzymes responsible for metabolism of LTC_4_ to LTD_4_ and LTE_4_, respectively, after its release from cells of which macrophages are the highest expressors (**Figure 3F**). Thus, GATA6 reduces expression of *Ltc_4_s* in correctly programmed pMФ reducing accumulation of LTE_4_ in the tissue. Somewhat surprisingly, while 5-HETE, LTB_4_ and LTC_4_ were included in our oxylipin panels, they were not reliably or consistently detected in lavage. This suggests that formation of LTE_4_ is fast, preventing its precursors from accumulating sufficiently to be detected.

### Eosinophils accumulate in homeostatic peritoneal cavity in *Gata6*-KO^mye^ microenvironment

Previously, we reported increased eosinophil numbers in the peritoneum of *Gata6*-KO^mye^ mice^6^. Here, we confirmed that the absolute number and proportion of Siglec-F^+^, SSC^high^ eosinophils in the peritoneal cavity of *Gata6*-KO^mye^ (where roughly one-fifth of total cells were found to be eosinophils) is increased (**Figure 4A and Figure S5A**). However, looking into the underlying mechanisms, in the bone marrow, where eosinophils differentiate, we found that their numbers are similar (**Figure S5B**) suggesting a local impact of *Gata6*-KO^mye^ in the peritoneal cavity. The impact of GATA6 deficiency was more pronounced in females and limited to peritoneal cavity only (**Figures 4A**, **S5B**). This high frequency of eosinophils was established in *Gata6*-KO^mye^ as early as 5 weeks of age. In contrast, in *Gata6*-WT, eosinophils expressed as a proportion of total cells, were highest at 2-3 weeks of age then dropped significantly afterwards (**Figure S5C**). The frequency of pMФ within the peritoneal cavity remained similar at different ages within the genotypes (**Figure S5D**). As females showed higher eosinophil levels, we next focused on this gender only for characterisation of their regulation.

**Figure 4:**
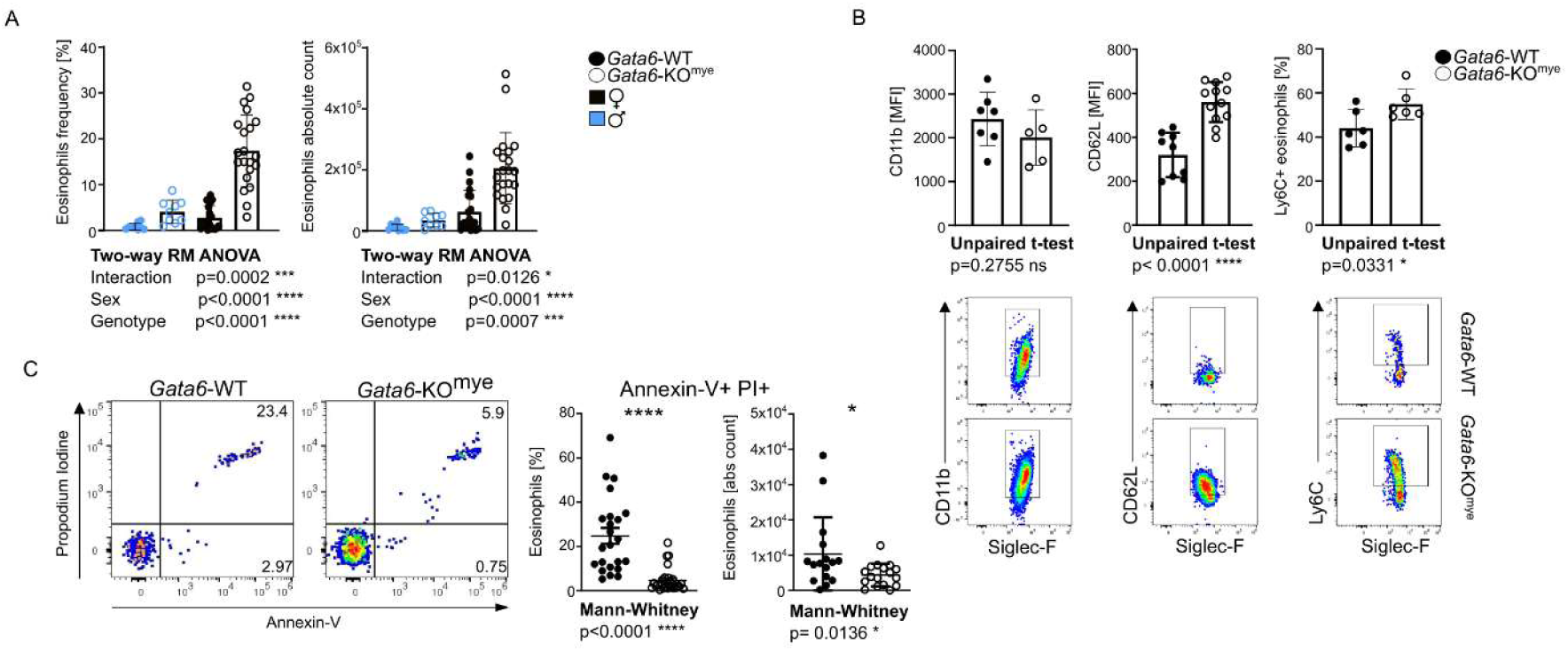
Dysregulated peritoneal eosinophil homeostasis in the absence of GATA6. **(A)** Eosinophil proportion and absolute count in the peritoneal cavity of *Gata6*-WT and *Gata6*-KO^mye^ female (black) and male (blue) mice. **(B)** Marker surface expression on peritoneal eosinophils (pre-gated by their characteristic SSC^high^, Siglec-F^+^ phenotype) in *Gata6*-WT and -KO^mye^ mice with corresponding representative flow cytometry density plots. **(C)** Annexin and PI staining of freshly isolated peritoneal eosinophils from *Gata6*-WT and *Gata6*-KO^mye^ mice and quantification of Annexin V and PI double positive eosinophils. Indicated statistical analysis was performed. *p<0.05, ***p<0.001, ****p<0.0001. Each symbol represents an individual mouse.

Eosinophilia is a feature of inflammation, associated with tissue damage caused by release of cell granules following activation or cell death^46^. Upregulation of Siglec-F and CD69, and downregulation of CD62L (L-selectin) are indicators of eosinophil activation in mice^47^. Investigation of eosinophil surface marker expression in *Gata6-*WT *and Gata6*-KO^mye^ peritoneal cavity, showed high levels of Siglec-F, CD11b and a characteristic flow-cytometric SSC^high^ profile^48^ consistent with their granularity (**Figure 4B and Figure S5A and E**). Using this characteristic expression profile to gate on eosinophils, we found that expression of all investigated eosinophil markers were unaffected (**Figure S5E**) apart from CD62L which was significantly higher on eosinophils from *Gata6*-KO^mye^ mice (p<0.0001) (**Figure 4B**). CD69 was not detected on eosinophils (data not shown). A subpopulation of Ly6C^+^ eosinophils was detected in both genotypes in line with previous studies^49^, and this was significantly increased in *Gata6*-KO^mye^ mice (**Figure 4B**). Collectively, the lack of CD69, no change in Siglec-F expression and upregulation of CD62L on the eosinophils that accumulate in *Gata6*-KO^mye^ peritoneal cavity indicates that they are no more activated than the wild type controls. Thus, observed accumulation of eosinophils in *Gata6*-KO^mye^ peritoneal cavity could be attributed to increased migration of these cells or prolonged survival in the tissue. To investigate these alternatives, we next investigated the levels of chemokines involved in eosinophil migration.

Eosinophils differentiate in the bone marrow and migrate in response to a gradient of eotaxin1 (CCL11) and eotaxin2 (CCL24) towards peripheral tissues, where they survive for several days^50^. To test whether eosinophilia in the *Gata6*-KO^mye^ peritoneal cavity is due to increased levels of these chemokines, CCL11 and CCL24 were measured in cell free peritoneal cavity lavages. CCL11 was unchanged in *Gata6*-KO^mye^, while CCL24 was significantly reduced (**Figure S5F**). Also, the amount of these chemokines did not correlate with eosinophil numbers (**Figure S5G**). These data suggest that CCL11/24 mediated migration of eosinophils to the peritoneal cavity of *Gata6*-KO^mye^ mice most likely does not account for eosinophilia observed.

Next, we investigated apoptosis of peritoneal eosinophils in *Gata6-*WT and *Gata6*-KO^mye^ mice, as a measure of clearance. Propidium Iodide/Annexin-V staining was reduced in peritoneal eosinophils from *Gata6*-KO^mye^ mice (**Figure 4C**). Decreased apoptosis of *Gata6*-KO^mye^ eosinophils was further verified using SYTOX AADvanced/Caspase 3/7 staining (**Figure S5H**). Thus, the accumulation of eosinophils in *Gata6*-KO^mye^ mice peritoneal cavity is associated with reduced cell death rather than increased migration to the tissue. To determine the signals prolonging eosinophil survival in *Gata6*-KO^mye^ we next focused on known survival signals for eosinophils and the disrupted lipidome described earlier.

### Lipidome changes in *Gata6*-KO^mye^ pMФ drive prolonged survival of eosinophil in the peritoneal cavity

Eosinophil survival is known to be regulated by several cytokines and lipid mediators. For example, IL-5 and granulocyte/macrophage-colony stimulating factor (GM-CSF) are recognised for their roles in maintaining eosinophil survival^51^. To identify what factors may regulate their survival here, we determined the levels of 40 cytokines and other mediators in cell free peritoneal lavage (**Figure S6A-D**). The overall cytokine/chemokine profile was similar for both genotypes with no significant detection of IL-5 and GM-CSF by protein array (**Figure S6A-D**). IL-5 was detected at low levels using ELISA, but this was unaffected by genotype (**Figure 5A**) and did not correlate with eosinophil numbers (**Figure S6E**). Considering the essential role of IL-5 in eosinophil migration and survival, *Gata6*-WT and *Gata6*-KO^mye^ mice were next challenged intraperitoneally with either α-IL-5 neutralising antibody (TRGK5) or a matched isotype control. At 72 hours post-injection, there was a significant drop in eosinophil numbers in the peritoneal cavity in both genotypes (**Figure 5B**). Additionally, in both, bone marrow and peripheral blood, IL-5 neutralization reduced eosinophil counts (**Figure 5C,D and S6F**). α-IL-5 treatment also significantly increased apoptosis of eosinophils in both genotypes (**Figure 5E**). Thus, our results demonstrate that IL-5 inhibition reduces eosinophil numbers in the bone marrow, blood, and peritoneal cavity, as well as promoting their cell death by apoptosis. Lastly, eosinophil apoptosis in α-IL-5 treated *Gata6*-WT peritoneum was significantly higher compared to *Gata6*-KO^mye^. Thus, whilst IL-5 is required generally for eosinophil survival in the peritoneal cavity in both genotypes there is no obvious differential effect. This indicates that other survival signals are responsible for the higher eosinophil counts seen in the peritoneal cavity of the *Gata6*-KO^mye^ mice when compared to the *Gata6*-WT mice.

**Figure 5:**
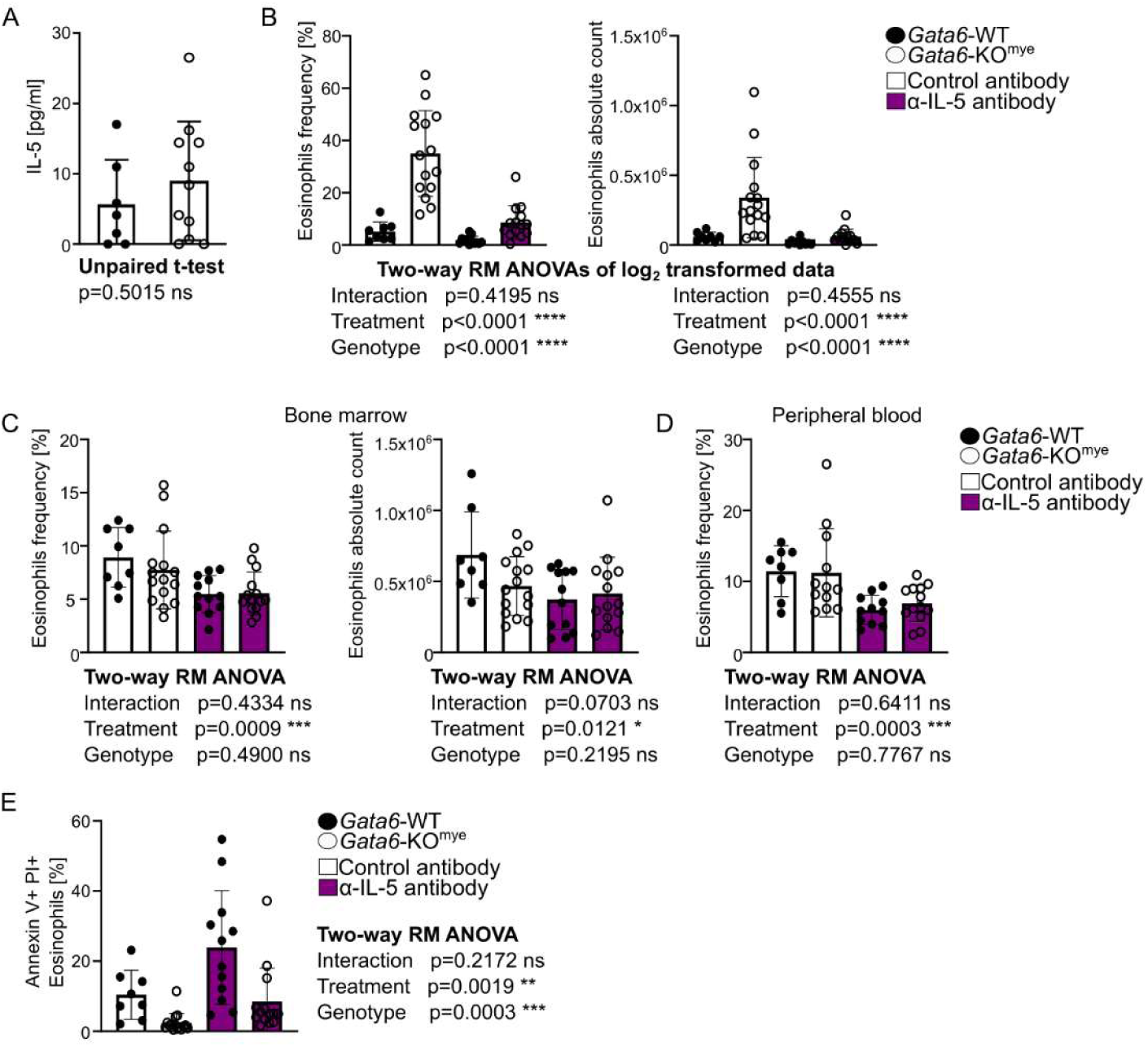
Role of IL5 in peritoneal eosinophil regulation. **(A)** Levels of IL5 (measured by ELISA) in cell free peritoneal lavage obtained from steady-state *Gata6*-WT and *Gata6*-KO^mye^ mice. **(B-D)** *Gata6*-WT and *Gata6*-KO^mye^ mice were treated with 20 µg of α-IL5 neutralising antibody (TRGK5) or isotype matched control antibody via intraperitoneal injection. After 72 hours, the proportion and absolute count of eosinophils in the **(B)** peritoneal cavity, **(C)** bone marrow and **(D)** peripheral blood was measured. **(E)** Proportion of Annexin V and propidium iodine (PI) double positive eosinophils in the peritoneal cavity was investigated in the same samples. Data are expressed as mean ± SD and analysis were performed using two-way ANOVA analysis, apart from (A) where t-test was used. Each symbol represents an individual mouse.

Oxylipins have been previously linked to eosinophil survival *in vitro*. For example, PGE_2_ prolongs eosinophil survival^52^, while cysteine leukotrienes have been reported to be either pro-survival^53, 54^ or anti-inflammatory with no impact on survival^55, 56^. Therefore, we tested whether endogenously-generated oxylipins contribute to the increased eosinophil survival observed *in vivo* in *Gata6*-KO^mye^. First, mice were orally administered indomethacin to inhibit generation of prostaglandins (**Figure 6A**). As expected, this caused a significant downregulation of COX-derived TxB_2_, 6-keto PGF-1α, PGE_2_ and 11-HETE (**Figure 6B and S7A**). Elevated levels of 12-HETE were detected, which may have resulted from greater availability of substrate for 12/15-LOX oxygenation (**Figure S7B**). Indomethacin significantly increased eosinophil accumulation and reduced their apoptosis in the peritoneal cavity of both *Gata6*-WT and *Gata6*-KO^mye^ mice (**Figure 6C**), however there was no difference in the response between the genotypes. This suggests an inhibitory effect of prostaglandins on this process or increased availability of the survival signal in the tissue upon COX pathway inhibition. Next, we evaluated the role of 12/15-LOX (*Alox15*) in eosinophil accumulation as both *Alox15* (**Figure S4D**) and 12-HETE lipid levels (**Figure 3B**) were found to be significantly downregulated in *Gata6*-KO^mye^. Homozygous, *Alox15^-/-^* mice, exhibited no significant difference in eosinophil frequencies in the peritoneal cavity compared to C57BL/6J controls (**Figure S7C**), indicating that 12/15-LOX-derived oxylipins are not major mediators of eosinophil accumulation. Together, these data suggest that oxylipins from 12/15-LOX or COX are not responsible for the higher levels of eosinophils in *Gata6*-KO^mye^.

**Figure 6:**
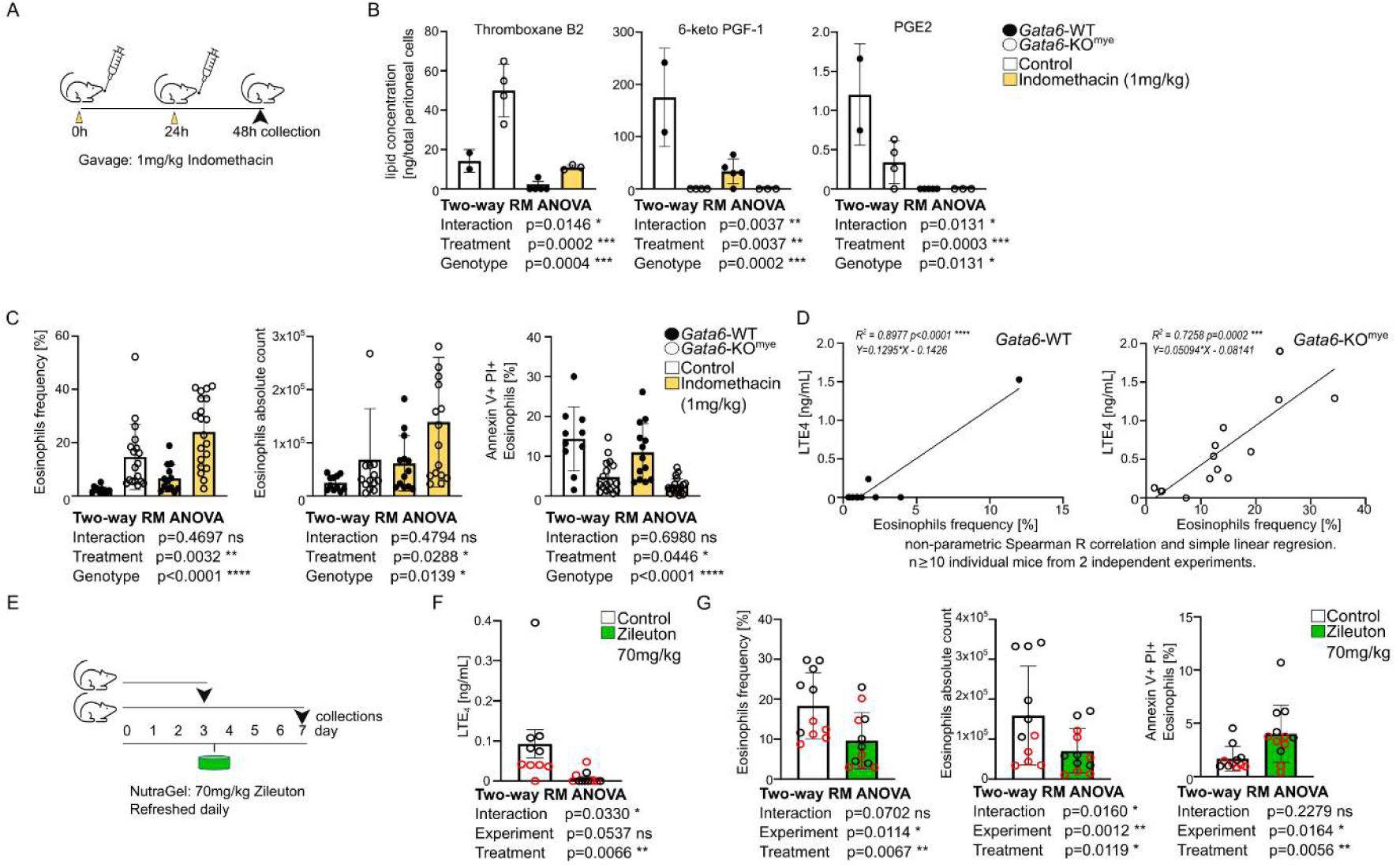
5-lipoxygenase inhibition counters increased eosinophil numbers seen in *Gata6*-KO^mye^ mice. **(A)** A schematic overview of the experiment performed in B and C. *Gata6*-WT and *Gata6*-KO^mye^ mice were gavaged with 1mg/kg of indomethacin or 0.05% Ethanol control, twice 24 hours apart. **(B)** Analysis of eicosanoids from a single experiment is shown *n≥2*. **(C)** Peritoneal lavage samples were collected from these mice 24 hours after the last treatment and were investigated for the proportion, absolute count and percentage apoptosis (Annexin V, propidium iodine) of eosinophils. *n≥10* individual mice per group, from 4 independent experiments. **(D)** Correlation plots between the amount of detectable LTE_4_ in cell free peritoneal lavage from *Gata6*-WT and *Gata6*-KO^mye^ mice and the frequency of eosinophils in these samples. Non-parametric Spearman R correlation and simple linear regression. n≥11 individual mice from 2 independent experiments. **(E)** A schematic overview of the experiment performed in F and G. *Gata6*-KO^mye^ mice were fed a complete nutrition Nutra-Gel containing 70mg/kg of Zileuton or control Nutra-Gel without any addition for 7 days with fresh supplemented Nutra-Gel provided daily. Peritoneal lavages were performed after 3 (Supplementary Figure 7E, F) or 7 days and **(F)** the levels of LTE_4_ in cell free lavage and **(G)** the proportion, absolute count and percent apoptosis (Annexin V, Propidium Iodine (PI)) of eosinophils were measured. *n≥11* individual mice per group, from 2 independent experiments indicated by the symbol colour. Data are expressed as mean±SD and analysis were performed using two-way ANOVA analysis. *p<0.05, **p<0.01, ***p<0.001, ****p<0.0001.

Next, oxylipin levels detected in cell free peritoneum were correlated with eosinophil frequency (**Figure S4B**) in both *Gata6*-WT and -KO^mye^ mice. This identified two candidate lipids, 9-HODE (**Figure S7D**) and the CysLT, LTE_4_ (**Figure 6D**). However, 9-HODE levels were not significantly different in cell free peritoneum or in pMФ, eosinophils or mast cells from *Gata6*-KO^mye^, in contrast to LTE_4_ which was significantly increased (**Figure 3B, S4A**). To test for a role for LTE_4_, this pathway was targeted *in vivo* in *Gata6*-KO^mye^ mice using a 5-LO (*Alox5*) inhibitor (**Figure 6E**). Zileuton administration in food significantly reduced peritoneal cell free lavage levels of LTE_4_ (**Figure 6F and S7E**). Additionally, Zileuton significantly decreased peritoneal eosinophil numbers after 7 days of treatment. This effect was attributed to increased apoptosis (AnnexinV+ PI+ staining) of the cells (**Figure 6G and Figure S7F**). In summary, our data identify LTE_4_ as a key regulator of eosinophil apoptosis in the peritoneal cavity, whose levels are restricted by pMФ tissue-specific programming as a result of GATA6 expression. Importantly, the data identify LTE_4_ elevation as responsible for the higher eosinophil survival observed in the absence of GATA6.

## Discussion

Here, disruption of MФ GATA6 resulted in significant changes to the lipidome, along with downstream functional consequences for immune homeostasis in the peritoneal cavity *in vivo*. Specifically, elevated sphingolipids, dysregulated oxylipins, and increased eosinophil accumulation associated with reduced apoptosis, was observed and found to be directly controlled by LTE_4_. Such an anti-apoptotic role for LTE_4_ was not previously described, demonstrating the importance of tissue-specific programming of the MФ lipidome in regulating local tissue immunity.

GATA6-deficiency represents a model of a tissue MФ lacking its normal specialisation. We and others previously demonstrated that this transcription factor is required by pMФ to deliver the characteristic homeostatic transcriptome of tissue resident macrophages in the peritoneum, including, regulation of renewal, survival and local immunity^5–7^. Our new data reveal that the loss of tissue-specialisation is associated with significant alterations to the cellular lipidome evidenced by accumulation of lipid-rich vesicles and alterations in lipid species in particular SL and oxylipins. Importantly, accumulation of SL is a characteristic feature of the sphingolipidoses, lysosomal storage disorders^14^. Thus, the lipid-rich vesicles identified in *Gata6*-KO^mye^ pMФ may represent uncatabolized SL in lysosomes. Consistent with SL accumulation, *Gata6*-KO^mye^ pMФ exhibited altered expression of their metabolic enzymes, including *Smpd1* and *Gba2*, mutations of which lead Niemann-Pick disease^57^ and Gaucher disease^58^, respectively. Here, inhibition of either *Smpd1* or *Gba2* in MФ demonstrated that lipidome changes comparable to those seen in *Gata6*-KO^mye^ pMФ, can be replicated by similar changes in gene expression in the metabolic pathways. Similarly, genes involved in eicosanoid metabolism, such as *Tbxas1, Ltc4s* and *Ptgis*, were significantly altered in *Gata6*-KO^mye^ pMФ, leading to corresponding changes in TxB_2_, LTE_4_, and 6-keto-PGF1α.

Recent studies have suggested potential functional links between SL and oxylipins. For example, Cer, C1P and LacCer, which are elevated in *Gata6*-KO^mye^ pMФ, can activate PLA_2_, while in mast cells, enzymes involved in generating sphingolipids (endoplasmic reticulum membrane protein ORM1-like protein 3 (ORMDL3) and serine palmitoyltransferase1 (SPT1)) can functionally associate with 5-LOX, leading to simultaneous elevations in both SL and LTs ^39–42^. This suggests that SL metabolism may directly regulate LTs, and in support of this idea, our study provides an additional mechanism through the upregulation of *Ltc4s* in pMФ. Whether this upregulation is dependent on SL specifically remains to be established.

Noting the elevated levels of LTE_4_ in lavage of *Gata6*-KO^mye^ mice, we next focused on the impact of pMФ lipidome changes on modulation of the tissue microenvironment. LTC_4_ is released from cells upon synthesis and further metabolised to LTD_4_ and LTE_4_ by enzymes on the same cell or other cells (transcellular biosynthesis). Analysis of Immgen data^45^ predicted that not only pMФ, but also eosinophils could be involved, since they express *Ltc4s*, *GGT* and *DPEP*. The significant correlation of eosinophil frequency with LTE_4_ levels suggests that these cells could either be involved in the production of LTE_4_, depend on it for survival, or both. Whether CysLTs regulate eosinophil biology is currently unclear although CysLTs are increased in asthma^59, 60^, suggesting involvement in a disease where these cells are elevated. Previous studies on human peripheral eosinophils has shown that LTB_4_ and two CysLTs (LTC_4_, D_4_), or signalling through the CYSLTR1 receptor (a receptor for LTC_4_, D_4_ and E_4_) can increase survival^53, 54^, while others found no effect of either LTB_4_ or LTE_4_ on eosinophil apoptosis^55, 56^. All these studies used peripheral blood eosinophils which may differ considerably from those localised to the peritoneal cavity, in a different environmental milieu. Here, we found that murine eosinophils accumulate, seemingly nonactivated, in the healthy peritoneal cavity of *Gata6*-KO^mye^ mice, exhibiting 5-LOX dependent reduced apoptosis. This is most likely regulated by LTE_4_ as this was the only lipid detected from that pathway and as mentioned, its concentration correlated with the presence of eosinophils. While, the exact and sensitive receptor of LTE_4_ remains to be identified, studies show some level of potency of LTE_4_ to known CysLTs receptors, which are G*i* protein-coupled receptors (GPCR), and eosinophils’ migration dependence on GPCR signalling^50, 61, 62^. Thus, suggesting the potential role of GPCR receptors in eosinophil phenotype observed here, although this remains to be determined. Our observations of increased survival are consistent with prior studies of peritoneal eosinophil turnover, which indicated prolonged survival is usually responsible for tissue persistence rather than cell migration in response to chemokines such as CCL11/24^50^.

Although IL-5 was also required to maintain eosinophil numbers in the peritoneum by promoting survival or regulating migration, there was no differential impact of genotype seen. Instead, our data reveal LTE_4_ to be an auxiliary survival signal acting synergistically with IL-5 to regulate eosinophil accumulation in a *Gata6*-dependent manner. This is analogous to the ability of LTC_4_ to promote eosinophil accumulation via collaboration with IL-33 on ILC2 cell cytokine production^63^. Similarly, LTD_4_ and GM-CSF synergise *in vitro* to directly enhance proliferation of eosinophil progenitors^64^. Thus, our observation that LTE_4_ can synergize with IL-5 extends our knowledge of oxylipins acting collaboratively with cytokines to regulate eosinophil biology *in vivo*.

Current therapies for eosinophilic diseases focus on depleting eosinophils, but their exact function is not fully understood, with emerging research highlighting their important antitumor, anti-inflammatory and pro-resolving roles in tissues^65–67^. Previously, it was proposed that LTE_4_ inhibits eosinophil degranulation^56^, making modulation of eosinophil activation an attractive therapeutic target. However, the molecular basis of this remains unclear. Our study reveals a tissue-specific programming of macrophages and their bioactive lipids limits eosinophil accumulation at steady-state and could provide an alternative approach for regulating eosinophil function and numbers in disease. While the 5-LOX inhibitor Zileuton is used in clinics for prevention of asthma attacks, whether it has any impact on eosinophil persistence is not yet known^68^.

In summary, our study advances the understanding of the role of tissue-resident macrophage programming in dictating the macrophage lipidome. Disruption of this programming, by GATA6 deletion in pMФ, revealed a homeostatic mechanism that restricts the availability of CysLT, limiting the accumulation of tissue eosinophils by regulating their survival. Our study demonstrates both the importance of tissue macrophage programming in regulating the cellular lipidome and highlights a mechanism by which macrophages can control the tissue microenvironment and the biology of co-localised cells, specifically survival and activation of tissue-associated eosinophils.

## Acknowledgements

We thank Dr John Watkins for the advice on the statistical analysis and Dr Brendon Naicker for help with untargeted LC-MS analysis. We thank Mark Gurney for technical support, and all animal facilities staff for help with colonies maintenance. For the purpose of open access, the author has applied a CC BY public copyright licence to any Author Accepted Manuscript version arising. This work was supported by Biotechnology and Biological Sciences Research Council Discovery Fellowship (BB/T009543/1) to MAC. P.R.R, J.A.J, C.H and V.O.D were supported by European Research Council (LIPID ARRAYS). P.R.T was funded by a Wellcome Trust Investigator Award (107964/Z/15/Z), an MRC Project Grant (MR/J002151/1), a UK Dementia Research Institute Programme (UK DRI-3001) and he is also supported by the Moondance Foundation.

## Authors Contributions

MAC, PRT and VBO’D conceived the project and designed the research; MAC, PR, NI, MR, SD, IP, CH, DF, JA-J, VJT, WL performed the experiments; MAC, PR, VJT, and VBO’D analysed mass spectrometry data; MAC, PR NI, RA and PRT examined biological data; IP and PB examined CARS data; MAC, PR, VBO’D and PRT wrote the manuscript.

## Conflict of Interest

The authors declare no conflicts of interest.

## Notes

### Competing Interest Statement

The authors have declared no competing interest.

## References

1. Yona, S. et al. Fate mapping reveals origins and dynamics of monocytes and tissue macrophages under homeostasis. Immunity 38, 79–91 (2013).

2. Ginhoux, F. & Guilliams, M. Tissue-Resident Macrophage Ontogeny and Homeostasis. Immunity 44, 439–449 (2016).

3. Guilliams, M., Thierry, G.R., Bonnardel, J. & Bajenoff, M. Establishment and Maintenance of the Macrophage Niche. Immunity 52, 434–451 (2020).

4. Davies, L.C., Jenkins, S.J., Allen, J.E. & Taylor, P.R. Tissue-resident macrophages. Nat Immunol 14, 986–995 (2013).

5. Gautier, E.L. et al. Gata6 regulates aspartoacylase expression in resident peritoneal macrophages and controls their survival. J Exp Med 211, 1525–1531 (2014).

6. Rosas, M. et al. The transcription factor Gata6 links tissue macrophage phenotype and proliferative renewal. Science 344, 645–648 (2014).

7. Okabe, Y. & Medzhitov, R. Tissue-specific signals control reversible program of localization and functional polarization of macrophages. Cell 157, 832–844 (2014).

8. Cassado Ados, A., D’Imperio Lima, M.R. & Bortoluci, K.R. Revisiting mouse peritoneal macrophages: heterogeneity, development, and function. Front Immunol 6, 225 (2015).

9. Wang, J. & Kubes, P. A Reservoir of Mature Cavity Macrophages that Can Rapidly Invade Visceral Organs to Affect Tissue Repair. Cell 165, 668–678 (2016).

10. Cantuti-Castelvetri, L. et al. Defective cholesterol clearance limits remyelination in the aged central nervous system. Science 359, 684–688 (2018).

11. Uderhardt, S. et al. 12/15-lipoxygenase orchestrates the clearance of apoptotic cells and maintains immunologic tolerance. Immunity 36, 834–846 (2012).

12. Ipseiz, N., et al. Tissue-resident macrophages actively suppress IL-1beta release via a reactive prostanoid/IL-10 pathway. EMBO J, e103454 (2020).

13. Smith, J.D. et al. Decreased atherosclerosis in mice deficient in both macrophage colony-stimulating factor (op) and apolipoprotein E. Proc Natl Acad Sci U S A 92, 8264–8268 (1995).

14. Schulze, H. & Sandhoff, K. Lysosomal lipid storage diseases. Cold Spring Harb Perspect Biol 3 (2011).

15. Misheva, M. et al. Oxylipin metabolism is controlled by mitochondrial beta-oxidation during bacterial inflammation. Nat Commun 13, 139 (2022).

16. Bligh, E.G. & Dyer, W.J. A rapid method of total lipid extraction and purification. Can J Biochem Physiol 37, 911–917 (1959).

17. Kessner, D., Chambers, M., Burke, R., Agus, D. & Mallick, P. ProteoWizard: open source software for rapid proteomics tools development. Bioinformatics 24, 2534–2536 (2008).

18. Dunn, W.B. et al. Metabolic profiling of serum using Ultra Performance Liquid Chromatography and the LTQ-Orbitrap mass spectrometry system. J Chromatogr B Analyt Technol Biomed Life Sci 871, 288–298 (2008).

19. Shaner, R.L. et al. Quantitative analysis of sphingolipids for lipidomics using triple quadrupole and quadrupole linear ion trap mass spectrometers. J Lipid Res 50, 1692–1707 (2009).

20. Gurney, M., et al. Lentiviral Vector Preparation for Efficient Gene and MicroRNA Modulation of Peritoneal Cavity Tissue-Resident Macrophages In Vivo in Mice. J Vis Exp (2024).

21. Ipseiz, N. et al. Effective In Vivo Gene Modification in Mouse Tissue-Resident Peritoneal Macrophages by Intraperitoneal Delivery of Lentiviral Vectors. Mol Ther Methods Clin Dev 16, 21–31 (2020).

22. Rosas, M., Thomas, B., Stacey, M., Gordon, S. & Taylor, P.R. The myeloid 7/4-antigen defines recently generated inflammatory macrophages and is synonymous with Ly-6B. J Leukoc Biol 88, 169–180 (2010).

23. Taylor, P.R. et al. The beta-glucan receptor, dectin-1, is predominantly expressed on the surface of cells of the monocyte/macrophage and neutrophil lineages. J Immunol 169, 3876–3882 (2002).

24. Davies, L.C. et al. A quantifiable proliferative burst of tissue macrophages restores homeostatic macrophage populations after acute inflammation. Eur J Immunol 41, 2155–2164 (2011).

25. Rosas, M. et al. The induction of inflammation by dectin-1 in vivo is dependent on myeloid cell programming and the progression of phagocytosis. J Immunol 181, 3549–3557 (2008).

26. Love, M.I., Huber, W. & Anders, S. Moderated estimation of fold change and dispersion for RNA-seq data with DESeq2. Genome Biol 15, 550 (2014).

27. Masia, F., Glen, A., Stephens, P., Borri, P. & Langbein, W. Quantitative chemical imaging and unsupervised analysis using hyperspectral coherent anti-Stokes Raman scattering microscopy. Anal Chem 85, 10820–10828 (2013).

28. Masia, F., Pope, I., Watson, P., Langbein, W. & Borri, P. Bessel-Beam Hyperspectral CARS Microscopy with Sparse Sampling: Enabling High-Content High-Throughput Label-Free Quantitative Chemical Imaging. Anal Chem 90, 3775–3785 (2018).

29. Karuna, A. et al. Label-Free Volumetric Quantitative Imaging of the Human Somatic Cell Division by Hyperspectral Coherent Anti-Stokes Raman Scattering. Anal Chem 91, 2813–2821 (2019).

30. Nahmad-Rohen, A. et al. Quantitative Label-Free Imaging of Lipid Domains in Single Bilayers by Hyperspectral Coherent Raman Scattering. Anal Chem 92, 14657–14666 (2020).

31. Di Napoli, C. et al. Hyperspectral and differential CARS microscopy for quantitative chemical imaging in human adipocytes. Biomed Opt Express 5, 1378–1390 (2014).

32. Di Napoli, C. et al. Quantitative Spatiotemporal Chemical Profiling of Individual Lipid Droplets by Hyperspectral CARS Microscopy in Living Human Adipose-Derived Stem Cells. Anal Chem 88, 3677–3685 (2016).

33. Pope, I. et al. Identifying subpopulations in multicellular systems by quantitative chemical imaging using label-free hyperspectral CARS microscopy. Analyst 146, 2277–2291 (2021).

34. Merrill, A.H. Sphingolipid Biosynthesis. In: Lennarz, W.J. & Lane, M.D. (eds). Encyclopedia of Biological Chemistry (Second Edition). Academic Press: Waltham, 2013, pp 281–286.

35. Eskes, E.C.B. et al. Biochemical and imaging parameters in acid sphingomyelinase deficiency: Potential utility as biomarkers. Mol Genet Metab 130, 16–26 (2020).

36. Schuchman, E.H. Acid sphingomyelinase, cell membranes and human disease: lessons from Niemann-Pick disease. FEBS Lett 584, 1895–1900 (2010).

37. Martin, E. et al. Loss of function of glucocerebrosidase GBA2 is responsible for motor neuron defects in hereditary spastic paraplegia. Am J Hum Genet 92, 238–244 (2013).

38. Woeste, M.A. & Wachten, D. The Enigmatic Role of GBA2 in Controlling Locomotor Function. Front Mol Neurosci 10, 386 (2017).

39. Bugajev, V. et al. Crosstalk between ORMDL3, serine palmitoyltransferase, and 5-lipoxygenase in the sphingolipid and eicosanoid metabolic pathways. J Lipid Res 62, 100121 (2021).

40. Pettus, B.J. et al. Ceramide 1-phosphate is a direct activator of cytosolic phospholipase A2. J Biol Chem 279, 11320–11326 (2004).

41. Nakamura, H. et al. Lactosylceramide-Induced Phosphorylation Signaling to Group IVA Phospholipase A(2) via Reactive Oxygen Species in Tumor Necrosis Factor-alpha-Treated Cells. J Cell Biochem 118, 4370–4382 (2017).

42. Huwiler, A., Johansen, B., Skarstad, A. & Pfeilschifter, J. Ceramide binds to the CaLB domain of cytosolic phospholipase A2 and facilitates its membrane docking and arachidonic acid release. FASEB J 15, 7–9 (2001).

43. Nakamura, H., Hirabayashi, T., Someya, A., Shimizu, M. & Murayama, T. Inhibition of arachidonic acid release and cytosolic phospholipase A2 alpha activity by D-erythro-sphingosine. Eur J Pharmacol 484, 9–17 (2004).

44. Dennis, E.A. & Norris, P.C. Eicosanoid storm in infection and inflammation. Nat Rev Immunol 15, 511–523 (2015).

45. Heng, T.S., Painter, M.W. & Immunological Genome Project, C. The Immunological Genome Project: networks of gene expression in immune cells. Nat Immunol 9, 1091–1094 (2008).

46. Spencer, L.A., Bonjour, K., Melo, R.C. & Weller, P.F. Eosinophil secretion of granule-derived cytokines. Front Immunol 5, 496 (2014).

47. Zhang, M. et al. Defining the in vivo function of Siglec-F, a CD33-related Siglec expressed on mouse eosinophils. Blood 109, 4280–4287 (2007).

48. Voehringer, D., van Rooijen, N. & Locksley, R.M. Eosinophils develop in distinct stages and are recruited to peripheral sites by alternatively activated macrophages. J Leukoc Biol 81, 1434–1444 (2007).

49. Percopo, C.M. et al. SiglecF+Gr1hi eosinophils are a distinct subpopulation within the lungs of allergen-challenged mice. J Leukoc Biol 101, 321–328 (2017).

50. Ohnmacht, C., Pullner, A., van Rooijen, N. & Voehringer, D. Analysis of eosinophil turnover in vivo reveals their active recruitment to and prolonged survival in the peritoneal cavity. J Immunol 179, 4766–4774 (2007).

51. Park, Y.M. & Bochner, B.S. Eosinophil survival and apoptosis in health and disease. Allergy Asthma Immunol Res 2, 87–101 (2010).

52. Peacock, C.D., Misso, N.L., Watkins, D.N. & Thompson, P.J. PGE 2 and dibutyryl cyclic adenosine monophosphate prolong eosinophil survival in vitro. J Allergy Clin Immunol 104, 153–162 (1999).

53. Lee, E., Robertson, T., Smith, J. & Kilfeather, S. Leukotriene receptor antagonists and synthesis inhibitors reverse survival in eosinophils of asthmatic individuals. Am J Respir Crit Care Med 161, 1881–1886 (2000).

54. Becler, K., Hakansson, L. & Rak, S. Treatment of asthmatic patients with a cysteinyl leukotriene receptor-1 antagonist montelukast (Singulair), decreases the eosinophil survival-enhancing activity produced by peripheral blood mononuclear leukocytes in vitro. Allergy 57, 1021–1028 (2002).

55. Murray, J. et al. Role of leukotrienes in the regulation of human granulocyte behaviour: dissociation between agonist-induced activation and retardation of apoptosis. Br J Pharmacol 139, 388–398 (2003).

56. Steinke, J.W., Negri, J., Payne, S.C. & Borish, L. Biological effects of leukotriene E4 on eosinophils. Prostaglandins Leukot Essent Fatty Acids 91, 105–110 (2014).

57. Schuchman, E.H. & Desnick, R.J. Types A and B Niemann-Pick disease. Mol Genet Metab 120, 27–33 (2017).

58. Sun, A. Lysosomal storage disease overview. Ann Transl Med 6, 476 (2018).

59. Vachier, I. et al. High levels of urinary leukotriene E4 excretion in steroid treated patients with severe asthma. Respir Med 97, 1225–1229 (2003).

60. Wang, H.B., Akuthota, P., Kanaoka, Y. & Weller, P.F. Airway eosinophil migration into lymph nodes in mice depends on leukotriene C(4). Allergy 72, 927–936 (2017).

61. Austen, K.F., Maekawa, A., Kanaoka, Y. & Boyce, J.A. The leukotriene E4 puzzle: finding the missing pieces and revealing the pathobiologic implications. J Allergy Clin Immunol 124, 406–414; quiz 415-406 (2009).

62. Lynch, K.R. et al. Characterization of the human cysteinyl leukotriene CysLT1 receptor. Nature 399, 789–793 (1999).

63. Lund, S.J. et al. Leukotriene C4 Potentiates IL-33-Induced Group 2 Innate Lymphoid Cell Activation and Lung Inflammation. J Immunol 199, 1096–1104 (2017).

64. Braccioni, F. et al. The effect of cysteinyl leukotrienes on growth of eosinophil progenitors from peripheral blood and bone marrow of atopic subjects. J Allergy Clin Immunol 110, 96–101 (2002).

65. Grisaru-Tal, S. et al. Metastasis-Entrained Eosinophils Enhance Lymphocyte-Mediated Antitumor Immunity. Cancer Res 81, 5555–5571 (2021).

66. Cosway, E.J. et al. Eosinophils are an essential element of a type 2 immune axis that controls thymus regeneration. Sci Immunol 7, eabn3286 (2022).

67. Goh, Y.P. et al. Eosinophils secrete IL-4 to facilitate liver regeneration. Proc Natl Acad Sci U S A 110, 9914–9919 (2013).

68. Berger, W., De Chandt, M.T. & Cairns, C.B. Zileuton: clinical implications of 5-Lipoxygenase inhibition in severe airway disease. Int J Clin Pract 61, 663–676 (2007).

69. Pope, I., Langbein, W., Watson, P. & Borri, P. Simultaneous hyperspectral differential-CARS, TPF and SHG microscopy with a single 5 fs Ti:Sa laser. Opt Express 21, 7096–7106 (2013).

70. Pluta, M. Nomarski’s DIC microscopy: a review, vol. 1846. SPIE, 1994.

71. Paar, M. et al. Remodeling of lipid droplets during lipolysis and growth in adipocytes. J Biol Chem 287, 11164–11173 (2012).

72. Li, S., Li, Y., Yi, R., Liu, L. & Qu, J. Coherent Anti-Stokes Raman Scattering Microscopy and Its Applications. Frontiers in Physics 8 (2020).

